# Time-dependent dual effect of NLRP3 inflammasome in brain ischemia

**DOI:** 10.1101/2020.10.08.332007

**Authors:** Alejandra Palomino-Antolin, Paloma Narros-Fernández, Víctor Farré-Alins, Javier Sevilla-Montero, Celine Decouty-Pérez, Ana Belen Lopez-Rodriguez, Nuria Fernández, Luis Monge, Ana I. Casas, María José Calzada, Javier Egea

**Author notes:** Corresponding author: Javier Egea: Research Unit, Hospital Santa Cristina. C/ Maestro Vives, 2-3. 28009. Madrid, Spain.

## Abstract

**Background and purpose:** Post-ischemic inflammation contributes to worsening of ischemic brain injury and in this process, the inflammasomes play a key role. Inflammasomes are cytosolic multiprotein complexes which upon assembly activate the maturation and secretion of the inflammatory cytokines IL-1β and IL-18. However, participation of the NLRP3 inflammasome in ischemic stroke remains controversial. Our aims were to determine the role of NLRP3 in ischemia and to explore the mechanism involved in the potential protective effect of the neurovascular unit.

**Methods:** WT and NLRP3 knock-out mice were subjected to ischemia by middle cerebral artery occlusion (60 minutes) with or without treatment with MCC950 at different time points post-stroke. Brain injury was measured histologically with 2,3,5-triphenyltetrazolium chloride (TTC) staining.

**Results:** We identified a time-dependent dual effect of NLRP3. While neither the pre-treatment with MCC950 nor the genetic approach (NLRP3 KO) proved to be neuroprotective, post-reperfusion treatment with MCC950 significantly reduced the infarct volume in a dose-dependent manner. Importantly, MCC950 improved the neuro-motor function and reduced the expression of different pro-inflammatory cytokines (IL-1β, TNF-α), NLRP3 inflammasome components (NLRP3, pro-caspase-1), protease expression (MMP9) and endothelial adhesion molecules (ICAM, VCAM). We observed a marked protection of the blood-brain barrier (BBB), which was also reflected in the recovery of the tight junctions proteins (ZO-1, Claudin-5). Additionally, MCC950 produced a reduction of the CCL2 chemokine in blood serum and in brain tissue, which lead to a reduction in the immune cell infiltration.

**Conclusions:** These findings suggest that post-reperfusion NLRP3 inhibition may be an effective acute therapy for protecting the blood-brain barrier in cerebral ischemia with potential clinical translation.

## Introduction

Cerebral ischemia is the fifth cause of death and first of adult disability worldwide. Ischemic stroke represents 85% of strokes, and currently, the only treatments are pharmacological intravenous recombinant tissue plasminogen activator (rtPA) injection within 4.5 h after stroke onset ^1^ and mechanical thrombectomy up to 24 h after stroke onset ^2^. However, only 10% of patients with ischemic stroke receive treatment with rtPA, due to numerous contraindications or exclusion criteria including intracranial hemorrhage, hypertension and significant head trauma or stroke within the previous 3 months ^3^. Ischemic stroke is a sudden interruption of cerebral blood flow in a particular brain region, leading to rapid neuronal death and the release of factors such as damage-associated molecular patterns (DAMPs). These factors initiate an inflammatory process in the surrounding area, which can increase oxidative stress, microvascular failure, blood-brain barrier (BBB) damage and brain edema.

Inflammation is a defensive host response required for resistance to infection, or to resolve tissue injury. When damage occurs at the central nervous system (CNS), microglia is rapidly activated, migrates to the focus of the lesion acquiring an activated phenotype characterized by the release of pro-inflammatory cytokines such as IL-1β, TNF-α or IL-6 ^4,5^. The pathological scenario of hyper-inflammation recruits different blood cells that infiltrate the damaged parenchyma for the resolution of inflammation. Evidence from preclinical models of stroke suggests that IL-1β contributes to a worsening of ischemic brain injury^6^.

Inflammasomes are cytosolic multiprotein complexes which, upon assembly, activate the pro-inflammatory caspase-1 that is responsible for the maturation and secretion of the inflammatory cytokines IL-1 β and IL-18, and additionally induce pyroptosis^7,8^. NLRP3, abbreviation for NACHT, LRR and PYD domains-containing protein 3, also known as cryopyrin, is the best studied inflammasome. NLRP3 inflammasome responds to a broad spectrum of activating stimuli that include a suite of bacterial, fungal and viral pathogen-associated molecular patterns (PAMPs) and damage-associate molecule patterns (DAMPs) such as ATP, uric acid crystals, crystalline and aggregated substances such as asbestos, silica, and amyloid-β fibrils ^9^. The involvement of the inflammasome in acute brain injuries, such as stroke or brain trauma, has been recently considered as an interesting scientific niche with a promising therapeutic potential. In fact, NLRC4 (NACHT, LRR, CARD domain-containing protein 4) and AIM2 (Absent In Melanoma 2) inflammasomes have been recently identified as suitable therapeutic targets for stroke^10^. However, the participation of the NLRP3 inflammasome in ischemic stroke remains controversial. Despite it has been shown that NLRP3 inflammasome does not participate in stroke ^10,11^, there are several recent publications suggesting that NLRP3 inhibition either improves ^12,13^ or worsens ^11^ cerebral ischemia/reperfusion. Here, we sought to contribute to the understanding of NLRP3 regulation and its important role in ischemic stroke using both genetic and pharmacological approaches to inhibit NLRP3. Our results demonstrate the key role of NLRP3 inflammasome activation after brain ischemia. Surprisingly, its activation reminds key right before ischemia onset being directly related to animal survival. However, NLRP3 inhibition during the first hours post-ischemia induced a marked protection of the neurovascular unit and may be a promising target in cerebral ischemia.

## Materials and Methods

### Study design

All animal experiments were performed after approval of the protocol by the Ethics Committee of Universidad Autónoma de Madrid (Madrid, Spain) according to the European Guidelines for the use and care of animals for research. All efforts were made to minimize animal suffering and to reduce the number of animals used in the experiments. Animals were housed under controlled conditions and allowed free access to water and standard laboratory chow. We used female and male C57Bl6/J and NLRP3-/- (KO) adult (3-4 months) and aged (12 months) 20-25 g or 25-30 g respectively, supplied by the animal facilities of Universidad Autónoma de Madrid (Madrid, Spain. IMSR Cat# JAX:000664, RRID: IMSR_JAX:000664). For the therapeutic window study, mice were treated with MCC950 (3 mg/kg) at different time points: before transient middle cerebral artery occlusion (tMCAO) (pretreated) or treated 1h and 2h post-reperfusion. For the dose response curve, mice were subjected to tMCAO and after cerebral ischemia, animals were randomly divided into the following experimental groups: vehicle-treated and 1, 3, 10 and 50 mg/kg MCC950-treated (50% female and 50% male in each group). At the end of the experiment, animals were euthanized by cervical dislocation.

### In vivo transient MCAO ischemia model

This model has previously been established in our group as described in Parada et al., 2019 for mice ^14^. C57Bl6/J mice were anesthetized with isoflurane (0.8% in oxygen), placed on a heating-pad, and corporal temperature was maintained at 37.0°C using a servo-controlled rectal probe-heating pad (Cibertec, Spain). Transient cerebral ischemia was induced using the intraluminal filament technique. Using a surgical microscope (Tecnoscopio OPMI pico, Carl Zeiss, Meditec Iberia SA, Spain), a midline neck incision was made and the right common and external carotid arteries were isolated and permanently ligated. A microvascular temporarily ligature was placed on the internal carotid artery to non-permanently block the blood flow. A silicon rubber-coated monofilament (6023910PK10, Doccol Corporation, Sharon, MA, USA) was inserted through a small incision into the common carotid artery and advanced into the internal carotid artery until resistance was felt. The tip of the monofilament was precisely located at the origin of the right middle cerebral artery and thus interrupting blood flow. The filament was held in place by a tourniquet suture in the common carotid artery to prevent filament relocation during the ischemia period. Animals were maintained under anesthesia during 1h occlusion followed by the reperfusion period which started when the monofilament was removed. Occlusion was confirmed by a laser measurement of blood flow (MOORFLPI-2), and animals showing less than 80% signal drop compared with baseline were excluded. After the surgery, wounds were carefully sutured and animals were allowed to recover from surgery in a temperature-controlled cage. Animals were provided with postoperative analgesia (0.5% lidocaine, subcutaneously, s.c.). Operation time per animal did not exceed 15 minutes. Animals were excluded from the stroke analysis if they died within the first 24 h, if an intracerebral haemorrhage occurred or in case the animal scored 0 on the Bederson score. Sham-operated animals underwent the same surgical procedure, but the monofilament was not advanced into the internal carotid artery.

### Neurological function assessment

The injury severity in all mice groups was assessed by Bederson Score, Elevated Body Swing Test and the Four-limbs Hanging Wire Test. The Bederson Score ^15^ is defined as follows: Score 0, no apparent neurological deficits; 1, body torsion and forelimb flexion; 2, right side weakness and thus decreased resistance to lateral push; 3, unidirectional circling behaviour; 4, longitudinal spinning or seizure activity; 5, no movement showed. Within the elevated body swing test, mice were held ∼1 cm from the tail base and then elevated above the surface in the vertical axis around 20 cm. A swing was considered whenever the animal moved its head out of the vertical axis to either the left or the right side (more than 10 degrees). Ratio of right/left swings were subsequently analyzed. Finally, to directly evaluate strength, the four-limbs hanging wire test was performed. The mouse was placed on the center of the wire with a diameter of 10 cm. Later, the wire was slowly inverted and placed at ∼40 cm above a paper towel bedding. The time until the mouse fell from the wire was recorded until a maximum time of 120 s.

### Two-dimensional laser speckle imaging techniques

Cortical cerebral blood flow (CBF) was monitored using the laser speckle technique. Briefly, a CCD camera Laser Speckle Contrast Imager (MOORFLPI-2) was positioned above the head and a laser diode (785 nm) illuminated the intact skull surface to allow penetration of the laser in a diffuse manner through the brain^16^. Speckle contrast defined as the ratio of the standard deviation of pixel intensity to the mean pixel intensity was used to measure CBF as it is derived from the speckle visibility relative to the velocity of the light-scattering particles (blood). This was then converted to correlation time values, which are inversely and linearly proportional to the mean blood velocity. Laser speckle perfusion images were obtained 10 min before tMCAO and continuing throughout the ischemic period until 5 min into the reperfusion. CBF was measured again in the same animals 24 hours after reperfusion.

### Measurement of infarct volume

Following euthanasia, brains were quickly removed and cut into four 2-mm thick coronal sections using a mouse brain slice matrix (Harvard Apparatus, Spain). Sections were stained for 15 min at room temperature with 2% 2,3,5-triphenyltetrazolium chloride (TTC; Sigma-Aldrich, The Netherlands) in PBS to visualize infarctions^17^. Indirect infarct volumes were calculated by volumetry (ImageJ software) according to the following equation: Vindirect (mm3) = Vinfarct x (1-(Vih – Vch)/Vch), where the term (Vih – Vch) represents the volume difference between the ischemic hemisphere and the control hemisphere and (Vih – Vch)/Vch expresses this difference as a percentage of the control hemisphere.

### Determination of blood-brain barrier disruption and brain edema

To determine the cerebral vasculature permeability and brain edema formation after 1h tMCAO, 2% Evans blue tracer (Sigma Aldrich, The Netherlands) was diluted in 0.9% NaCl and then injected intraperitoneally (i.p.) 1 hour after removing the filament. Slices were scanned and total Evans Blue extravasation area was measured in each slice using ImageJ.

### Determination of chemokine CCL2 levels in blood serum and brain tissue by ELISA

Blood serum and brain tissue were obtained from all mice group 24 hours after stroke. Serum was separated by centrifugation at 2500 rpm for 10 min at 4 °C. Brain tissue was mechanically homogenized in cold PBS and centrifuged at 1200 rpm for 10 min at 4 °C to collect the supernatant. The protein concentration of the supernatant was determined using BCA protein assay kit (Thermo Scientific, USA) and 100 µg were loaded to measure CCL2 levels by a specific ELISA kit (R&D Systems).

### Cerebral microvessel enrichment

Cerebral microvessels were isolated from the ischemic hemispheres of mice that had undergone tMCAO surgery. Briefly, the hemispheric brain tissue was minced and homogenized in disaggregation buffer (15 mM HEPES, 150 mM NaCl, 4 mM KCl, 3 mM CaCl2, 12 mM MgCl2, 0.5% BSA and protease inhibitors). The homogenate was centrifuged at 3200 rpm for 10 minutes. The pellet was resuspended in 20% dextran and centrifuged for 30 minutes at 3000 rpm and 4°C. The pellet was resuspended in PBS and filtered through a 100-μm nylon cell strainer (Corning® cell strainer, USA), followed by a new filtration through a 20-μm nylon mesh (Corning® cell strainer, USA). Retained microvessels on the mesh were washed with RIPA buffer (Abcam, UK) and stored until further analysis. Protein concentration was determined using BCA protein assay kit (Thermo Scientific, USA) and 40 μg of lysate were loaded in 15% acrylamide gels. Anti-glial fibrillary acidic protein (GFAP) (1:1000; Abcam, UK) was used as a control antibody for validation of succeeded vessel enrichment (Supplemental fig. 1).

### Quantitative Real-time PCR

Total RNA was extracted from brain tissue and vessel enrichment by TRIzol method (10296-028, Invitrogen, Carlsbad, CA, USA) and cDNA was synthesized using iScript cDNA Synthesis Kit (Biorad) following the manufacturer’s instructions. Quantitative polymerase chain reaction (qPCR) was performed using Power SYBR Green PCR Master Mix (Thermo Fisher) in 384-well format using a QuantStudio 5 PCR system (Applied Biosystems, USA). Data were normalized to the expression of the housekeeping gene B2M. Specific primers were designed using the NCBI nucleotide data base and Primer 3 software (http://biotools.umassmed.edu/bioapps/primer3_www.cgi). Primer sequences (purchased from Sigma or Metabion internacional AG) were checked with BLAST (http://blast.ncbi.nlm.nih.gov/Blast.cgi). The comparative CT method (or the 2ΔCT method) was used to determine differences in gene expression between B2M and control samples. The sequences of the primers used in the study are listed in Table 1.

**Table 1.**
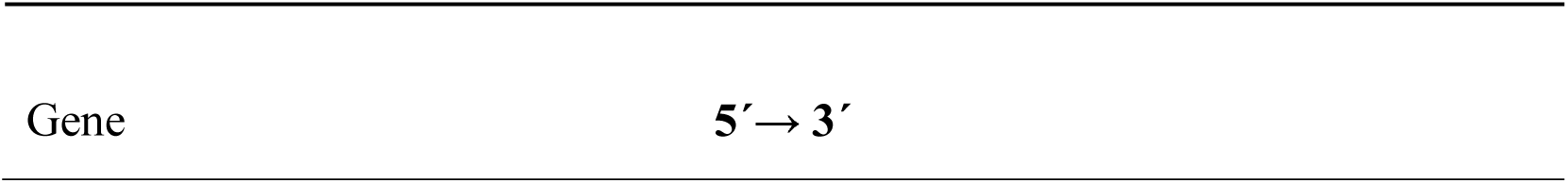
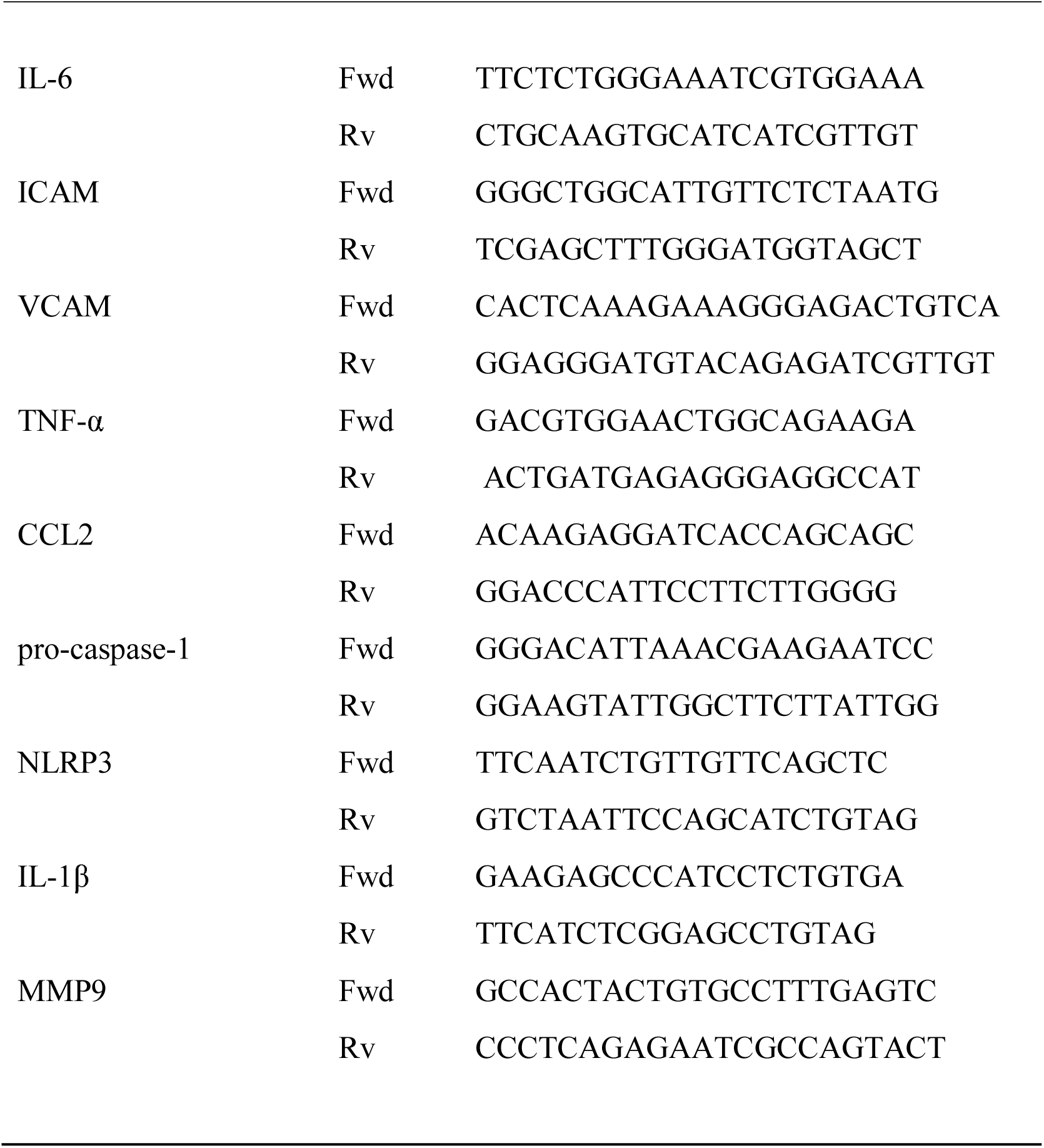
Primers usedin this study

### Immunohistochemistry. Immunofluorescence microscopy

After 24 hours of stroke, mice were perfused with PBS and brain were excised and freezing in an embedding optimal cutting temperature [OCT] compound on dry ice. Coronal slices of frozen brains were cut with a cryostat (CM 1100; Leica) at 10 μm thickness. Brain tissue cryosections were fixed with 4% PFA for 10 minutes at room temperature, washed with PBS, permeabilized in 0.3% Triton X-100/PBS for 15 min at room temperature, washed with PBS and blocked with 20% BSA over night at 4°C. Primary antibodies were diluted in PBS, BSA 1 % and incubated overnight at 4°C at the following dilutions: anti-ZO-1 (1:100, Zymed), anti-CD31 (1:100, BD Biosciences) anti-Claudin-5 (1:100, Abcam) and Ly6G (1:100, Abcam). After gentle washing, the sections were incubated with fluorochrome-conjugated secondary antibodies for 1 hour at room temperature (1:500 Alexa Invitrogen) and fluorescent DAPI (1:1000; ThermoFisher Scientific). Sections were washed in PBS and then mounted using a fluorescence mounting medium Fluoromount-G (SouthernBiotech, USA). Immunofluorescence images were obtained using a Leica TCS-SP5 (Leica Microsystems, Madrid, Spain) confocal microscope. Images were processed with the program ImageJ 1.52e (ImageJ software, National Institutes of Health, USA).

### Immunoblotting and image analysis

24 hours after stroke, brain tissue was collected and lysed in lysis buffer (1% Nonidet P-40, 10% glycerol, 137 mM NaCl, 20 mM Tris–HCl, pH 7.5, 1 g/mL leupeptin, 1 mM PMSF, 20 mM NaF, 1 mM sodium pyrophosphate, and 1 mM Na3VO4). Proteins (30 μg) from the cell lysate were resolved by SDS–PAGE and transferred to Immobilon-P membranes (Millipore Corp., Billerica, MA, USA). Membranes were incubated with anti-NLRP3 (1:1000; AdipoGene), anti-pro IL-1β (1:1000; R&D Systems), anti-Claudin-5 (1:500, Abcam), anti-ZO-1 (1:1000, Zymed), anti-β-actin (1:50000; Sigma-Aldrich) or anti-MMP9 (1:1000, Santa Cruz Biotechnology). Appropriate peroxidase-conjugated secondary antibodies (1:5000) were used to detect proteins by enhanced chemiluminescence. Different band intensities corresponding to immunoblot detection of protein samples were quantified using the Scion Image program (RRID:SCR_008673).

### Dissociation of cerebral tissue into single cell suspensions

Mice were euthanized by cervical dislocation and the brain was quickly and gently removed. The ischemic brain tissue (ipsilateral hemisphere) and the corresponding mirror regions of the non-ischemic tissue (contralateral hemisphere) were dissected out and analyzed separately. Ipsilateral and contralateral brain tissues were placed in cold Hank balanced salt solution (HBSS + Ca/Mg; Lonza) and mechanically dissected through a 100 µm cell strainer. Tissue suspension was centrifuged at 286 g for 5min at 4°C. Pellet was enzymatically digested in collagenase/liberase TL (2 U/mL) (Roche Diagnostics) for 1 h at 37 °C. The cell suspension was filtered through a 70 µm filter with DNAse (66 U/mL) (Roche Diagnostics). Cell pellet was resuspended in 25% density gradient and centrifuged at 521 g for 20 min at 18° C. The myelin coat was aspirated out and cells were collected from the interphase, washed once with HBSS and processed for flow cytometry.

### Flow cytometry

Isolated brain cells were washed with fluorescence-activated cell sorting (FACS) buffer (phosphate-buffered saline, 2 mM EDTA, 2% FBS), incubated at 4°C for 10 min with mouse FcBlock (BD Bioscience), and then with primary antibody in FACS buffer for 30 min at 4°C. The primary antibodies used were CD45 (Clone 30-F11, PB, 1:100, BD Pharmingen), CD11b (Clone M1/70, PE, 1:50, Miltenyi Biotec), Ly6G (clone 1A8, FITC, 1:100, BD Pharmingen), and Ghost viability dye (APC-H7, 1:100, Tombo Biosciences). Data were acquired on a BD FacsCanto II cytometer (BD Biosciences) using the FACS Diva software (BD Biosciences). Cells were morphologically identified by linear forward scatter (FSC-A) and side scatter (SSC-A) parameters. Data analysis was carried out with FlowJo (version 10.6.1. TreeStar Inc., Ashland, OR, USA). No less than 100,000 events were recorded for each sample. Cell type-matched fluorescence minus one (FMO) controls were used to determine the positivity of each antibody. The gating strategy employed to quantify frequencies of infiltrating and resident immune cells is shown in Fig. S2. Alive cells were identified based on viability dye fluorescence levels. Overall immune cells were identified based on expression of the leukocyte common antigen CD45 and additionally gated by CD11b expression to distinguish microglia (CD11bHiCD45Int) from non-myeloid (CD11b-CD45Hi) ^18^ and myeloid (CD11bHiCD45Hi) infiltrating immune cells ^19^. The myeloid subpopulation (CD11b+CD45Hi) was further gated to quantify neutrophils (Ly6GHi) ^20^. The frequencies of all immune cells were calculated from the alive cells.

### Statistical Analysis

Mice were randomly assigned to each experiment and treatment group and the blind analysis technique was used for all data. First, data were tested with Shapiro-Wilk test for normality and Levene’s test for equality of variance. Sex differences in infarct size, brain edema and neuro-motor functional tests were analyzed by two-way ANOVA. Since no sex difference was observed in any of these experiments, data from both male and female mice were pooled and analyzed together. All results were expressed as mean ± SD except for ordinal functional outcome scales that were depicted as scatter plots. Two-tailed unpaired Student t-test (Gaussian distribution) was used to analyze significant differences between 2 groups. Ordinary one-way ANOVA with Newman-Keuls Multiple Comparison post-test (Gaussian distribution) was used to compare variables among three or more groups. The number of animals necessary to detect a standardized effect size on infarct volumes ≥ 0.2 (vehicle treated control mice vs treated mice) was determined via a priori sample size calculation with the following assumptions: α = 0.05, ß = 0.2, mean, 20% SD of the mean. The threshold for statistical significance was p<0.05 throughout. All results were analyzed using the GraphPad Prism 6.0 software (GraphPad Software Inc., San Diego, CA, USA. RRID:SCR_008). All measurements were undertaken in at least three technical replicates.

## Results

### Post-reperfusion inhibition of NLRP3 inflammasome with MCC950 reduces infarct volume and neurological outcome

The involvement of the inflammasomes in stroke has been recently considered as an interesting scientific niche with a promising therapeutic potential. However, the participation of the NLRP3 inflammasome in ischemic stroke remains controversial so far. To evaluate the role of NLRP3 inflammasome in brain ischemia, wild type (WT) and NLRP3 -/- (KO) mice were subjected to the transient middle cerebral artery occlusion (tMCAO) model of brain ischemia (Fig.1A, top). To avoid any potential confounding effects, cerebral blood flow (CBF) was monitored by a Laser Speckle Contrast Imager (MOORFLPI-2) to ensure a complete cerebral flow blockage (Fig. 1A, bottom). To measure infarct volume, brain sections were stained 24 h post-reperfusion with 2,3,5-triphenyltetrazolium chloride (TTC) as described in ^17^. First, we evaluated the infarct volume in WT male and female mice (3-6 months), where no gender differences were identified (Fig. S3). Despite the infarct volume remained comparable in NLRP3 KO and vehicle WT mice (Fig. 1B), we identified a significant increase in the mortality rate of NLRP3 KO compared to non-genetically modified mice (50% vs 15.8%) (Table S1), suggesting a crucial role of NLRP3 activation in brain ischemia development and survival post-stroke. To evaluate a potential time-dependent role of NLRP3, both male and female C57Bl/6J mice were subjected to tMCAO and later treated with MCC950 (3 mg/kg) at different time points: (i) 30 min before tMCAO (pre-treatment), (ii) 1h and (iii) 2h post-reperfusion. Brain sections were stained with TTC 24h post-reperfusion to assess infarct volume (Fig. 1B). The pre-treatment with MCC950 30 min before tMCAO did not reduce infarct size but, similarly to the NLRP3 KO approach, mortality rate was increased (15.8% vs 63.6%) (Table S1). However, MCC950 (3mg/kg) treatment 1 h post-reperfusion significantly reduced infarct volume (113 ± 6 to 43 ± 9 mm3, p<0.01) while non-significant reduction was detected in treated animals 2h post-occlusion (113 ± 6 to 91 ± 14 mm3). Thus, these data confirm the dual role of NLRP3 activation in brain ischemia, being essential for survival post-stroke in a time-dependent manner. Hence, selection of the most suitable post-reperfusion time-point is crucial for defining the best pharmacological treatment using NLRP3 inhibitors.

**Figure 1.**
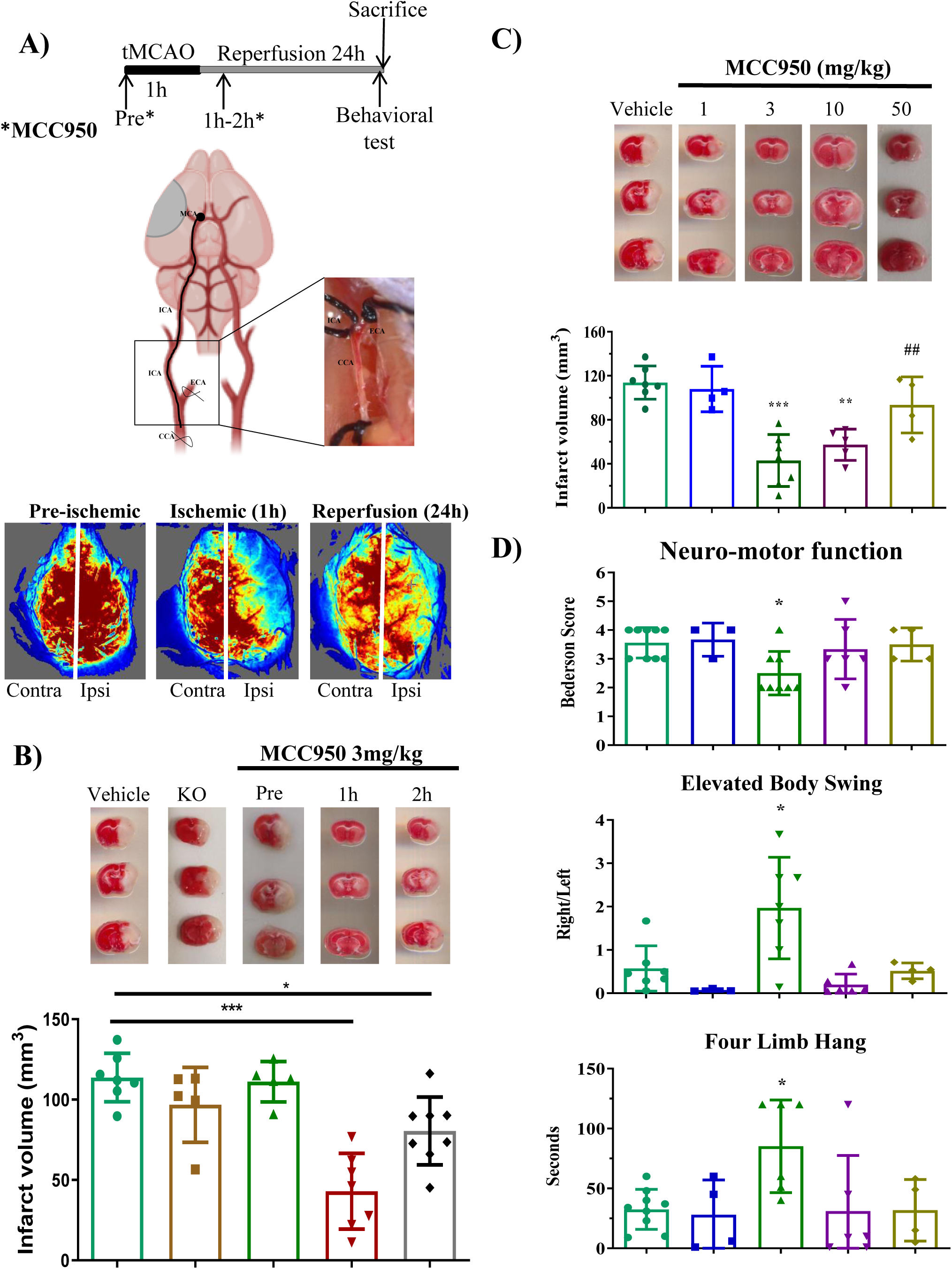
Effect of NLPR3 Inflammasome in cerebral ischemia 24 hours after 1 hour of stroke. (A) Adult mice were subjected to a 1h-transient occlusion of the middle cerebral artery (tMCAO) followed by 24h of reperfusion, before sacrificing. Treatment was injected i.p. pre-occlusion, 1h and 2h after the beginning of reperfusion. Representative images of cerebral blood flow before MCAO, during tMCAO, and at 10 min after reperfusion for each group. (B) Representative images and quantification of infarct volume by TTC staining at 24 hours after 1 hour of ischemia in NLRP3 -/- (n=5) and wild type adult mice treated 30 min pre-occlusion (n=5), 1h (n=7) and 2h (n=6) after reperfusion with MCC950 3mg/kg. (C) Representative images and quantification of infarct volume by TTC staining at 24hours after 1 hour of ischemia in adult mice treated 1h after reperfusion with Vehicle (n=7), MCC950 1mg/kg (n=4); 3mg/kg (n=7); 10mg/kg (n=5); 50mg/kg (n=4). (D) Neurological outcome quantification by Bederson Score, elevated body swing test and four limb hanging 24 hours after 1 hour of ischemia in adult mice treated with Vehicle (n=9), MCC950 1mg/kg (n=4); 3mg/kg (n=8); 10mg/kg (n=6); 50mg/kg (n=4). Data are mean ± SD. *p<0.05; **p<0.01; ***p<0.001 compared to vehicle mice; ##p<0.01 compared to MCC950 3 mg/kg.

Once we selected the best post-reperfusion time-point of MCC950 treatment, we performed a dose-response curve of MCC950 (1, 3, 10, 50 mg/kg) to choose the most effective dose. A significant reduction of brain infarct volume was shown in mice treated with MCC950 at 3 and 10 mg/kg compared with vehicle group (Fig. 1C). However, this protective effect was lost in animals treated with MCC950 at 50 mg/kg, which resulted in an increased mortality rate (50% vs 15.8%) (Table S1). Neurofunctional outcome and life quality are key parameters to be assessed in the clinical field. Thus, we simultaneously performed three independent neuromotor tests: Bederson Score, Elevated Body Swing test and the Four-Limb Hanging Wire test (Fig. 1D). All three tests were significantly improved in mice treated with MCC950 (3 mg/kg) 1 h after reperfusion, while treated animals at 1, 10 or 50 mg/kg at the same time point, showed no significant neuromotor improvement. Finally, we observed a significant reduction of infarct volume in middle-aged mice (12 months) treated with MCC950 (3 mg/kg) in comparison with vehicle-treated aged mice. However, we did not notice an improvement of the neurofunctional outcome in aged mice treated with MCC950 (3 mg/kg) (Fig. S4). Therefore, we established that the best conditions for NLRP3 inhibition with MCC950 are 3 mg/kg 1 h post-reperfusion, which reduced infarct volume and lead to neuromotor functioning amelioration in adult mice (3-4 months).

### MCC950 treatment prevents stroke-induced NLRP3 inflammasome activation

NLRP3 inflammasome is an important sensor that triggers the inflammatory response leading to the maturation and secretion of the inflammatory cytokines IL-1β and IL-18 ^21^. To determine the mechanistic NLRP3 inflammasome activation in the penumbra region, we analyzed the main components of NLRP3 inflammasome assembly (NLRP3 sensor, pro-caspase-1 and pro-IL-1β) 24 hours after stroke (Fig. 2A). NLRP3 and pro-caspase 1 mRNA levels were increased in vehicle tMCAO group compared with the sham group (20- and 25-fold, respectively). A significant decrease in mRNA levels was observed in animals treated with MCC950 (3mg/kg) (Fig 2B). In line with mRNA levels, NLRP3 protein expression was increased in non-treated stroked animals compared with the sham group. Nevertheless, treatment with MCC950 (3mg/kg) did not provide a marked reduction on NLRP3 protein levels. Indeed, we measured protein levels of pro-IL-1β and its active form IL-1β as an indicative of NLRP3 inflammasome activation. Moreover, while the levels of pro-IL-1β remained unaltered among the different groups, cleaved IL-1β protein levels were higher in the ischemic group compared to sham, and the treatment with MCC950 (3mg/kg) prevented the cleavage of IL-1β to its active form (Fig. 2C). These results demonstrate that MCC950 at 3 mg/kg effectively inhibits the activation of NLRP3 inflammasome and IL-1β cleavage after 24 hour of brain ischemia.

**Figure 2.**
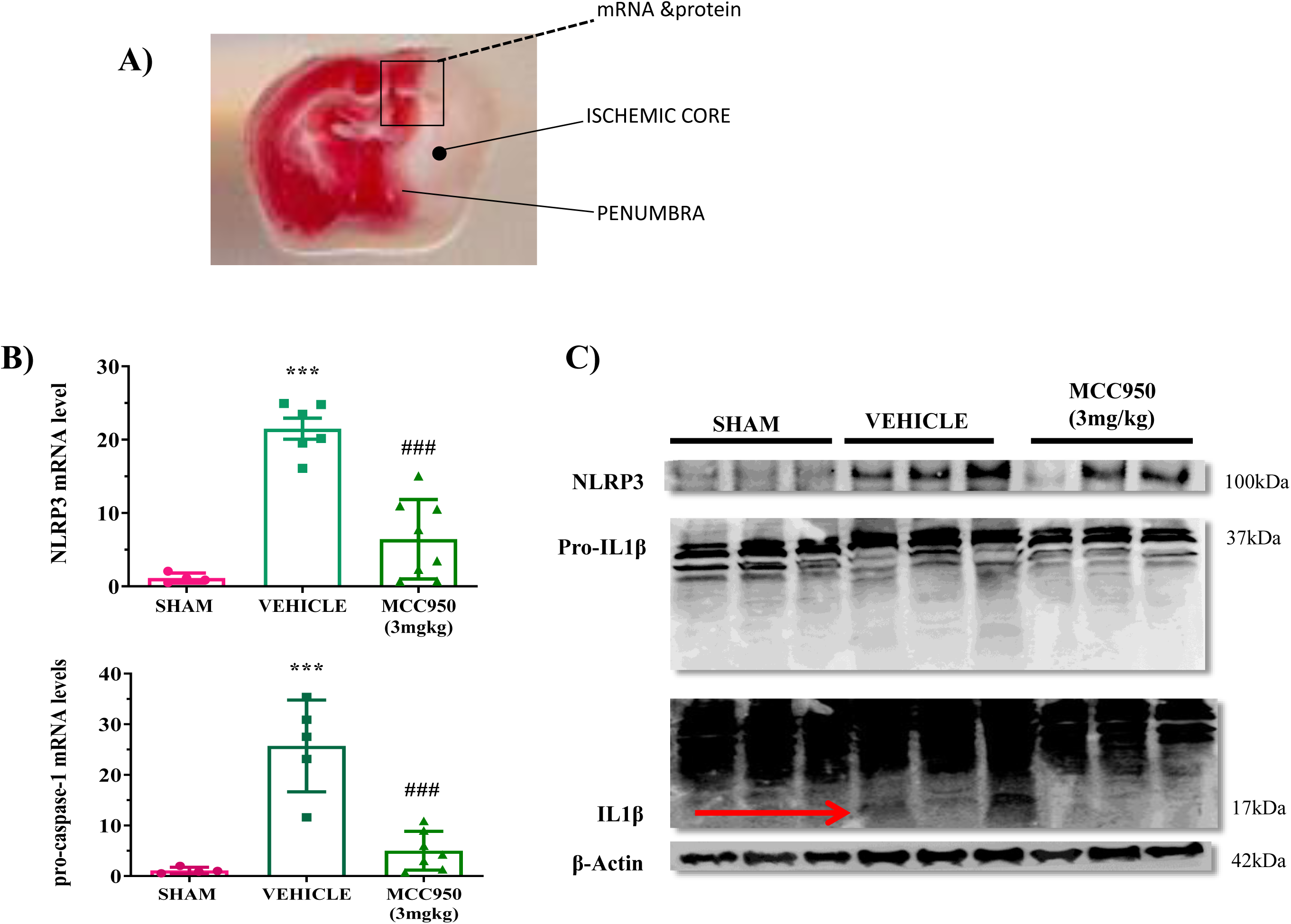
MCC950 at 3mg/kg inhibits NLRP3 inflammasome activation after brain ischemia. (A) Within the ischemic brain, there are two major areas of injury: the core ischemic zone and the ischemic penumbra. For further analysis we extracted tissue from both areas. (B) mRNA expression levels of the NLRP3 and pro-caspase-1 24h after 1 hour of stroke. (C) Western blot images show the expression of NLRP3, inactive pro-IL-1β and active IL-1β 24h after 1 hour of stroke in the brains of indicated groups of mice receiving sham plus vehicle, MCAO plus vehicle, or MCAO plus MCC950 at 3mg/kg. Data are mean ± SD. ***p<0.001 compared to sham mice; ###p<0.001 compared to vehicle mice.

### Pharmacological inhibition of inflammasome NLRP3 reduces the “cytokine storm” after brain ischemia

Immunity and inflammation are integral components of the pathogenic processes triggered by Ischemia-Reperfusion (I/R). Inflammatory signaling is responsible for early molecular events caused by I/R leading to brain invasion by blood-borne leukocytes ^22^. Hence, we examined whether NLRP3 inhibition is involved in inflammatory response after brain ischemia. We evaluated mRNA levels of three different pro-inflammatory cytokines (TNF-α, IL-6 and IL-1β), as well as the chemokine CCL2. MCC950 (3 mg/kg) significantly reduced IL-1β, TNF-α, and CCL2 levels in treated mice 1 h post-stroke (Fig. 3A, B, D). However, MCC950 treatment did not affect IL-6 mRNA levels (Fig. 3C). High levels of CCL2 chemokine are found in the brain perivascular space in many CNS pathologic states accompanied by inflammation and blood-brain barrier (BBB) disruption ^23^. To evaluate the effect of NLRP3 inhibition on the levels of CCL2 chemokine into the blood-serum space and the brain parenchyma after ischemia, we measured CCL2 in brain homogenates of sham, vehicle- and MC950-treated mice 24h post-reperfusion. We detected a significant increase of CCL2 in the ipsilateral hemisphere of vehicle-treated mice (82 ± 16 pg/mg), which was significantly reduced in the ipsilateral hemisphere of MCC950-treated mice (40 ± 11 pg/mg). As an internal control, no changes were found in the contralateral hemisphere of MCC950- or vehicle-treated mice (Fig. 3E). Moreover, we detected an increase of serum levels CCL2 in non-treated mice that were significantly decreased in mice subjected to MCC950 (Fig. 3F). Altogether, we here demonstrate that MCC950 treatment 1 hour after brain ischemia significantly reduces the inflammatory cytokine storm.

**Figure 3.**
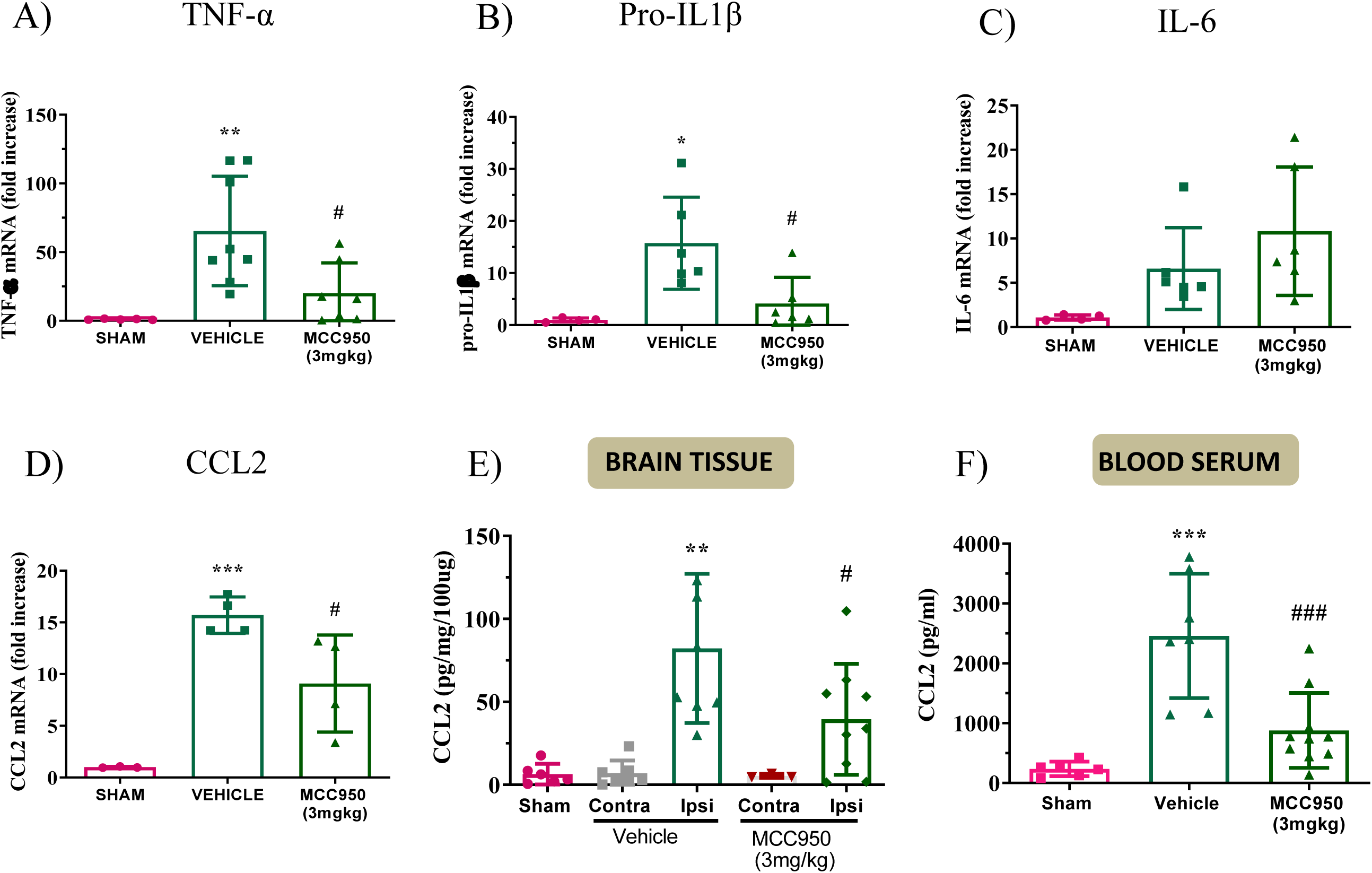
Inhibition of NLRP3 inflammasome affects the gene expression of different pro-inflammatory cytokines (TNF-α, pro-IL-1β and IL-6) and the level of chemokine CCL2 in brain tissue and blood serum after 24 hours of brain ischemia. (A, B, C) Cytokines (TNF-α, pro-IL-1β and IL-6) and chemokine (CCL2) (D) mRNA levels after 24 hours of brain ischemia in mice treated with MCC950 and vehicle. Data are normalized to sham. (E) Release of CCL2 in cerebral parenchyma after 24 hours of brain ischemia in mice with vehicle, and MCC950 treatment in contralateral and ipsilateral hemisphere. Data are normalized to sham. (F) Release of CCL2 in blood serum after 24 hour of brain ischemia with vehicle and MCC950 treatment. Data are normalized to sham. Data are mean ± SD. *p<0.05; **p<0.01; ***p<0.001 compared to sham mice; #p<0.05 and ###p<0.001 compared to vehicle mice.

### NLRP3 inflammasome inhibition preserves blood-brain barrier (BBB) integrity after brain ischemia

Under ischemic conditions, the BBB is disrupted followed by extravasation of blood components into the brain and therefore compromising the neuronal function. As CCL2 mediates endothelial dysfunction that may contribute to BBB disruption, we explored the potential effect of MCC950 treatment on the BBB disruption produced in the ischemic brain. Hence, we first analyzed BBB integrity after ischemic stroke by measuring Evans Blue extravasation, a vascular tracer, into the brain parenchyma. MCC950 (3 mg/kg) treatment significantly reduced the extravasation of Evans blue in stroke mice after 24 h compared with vehicle-treated mice (Fig. 4B), suggesting that NLRP3 play a key role in microvasculature stability post-stroke.

**Figure 4.**
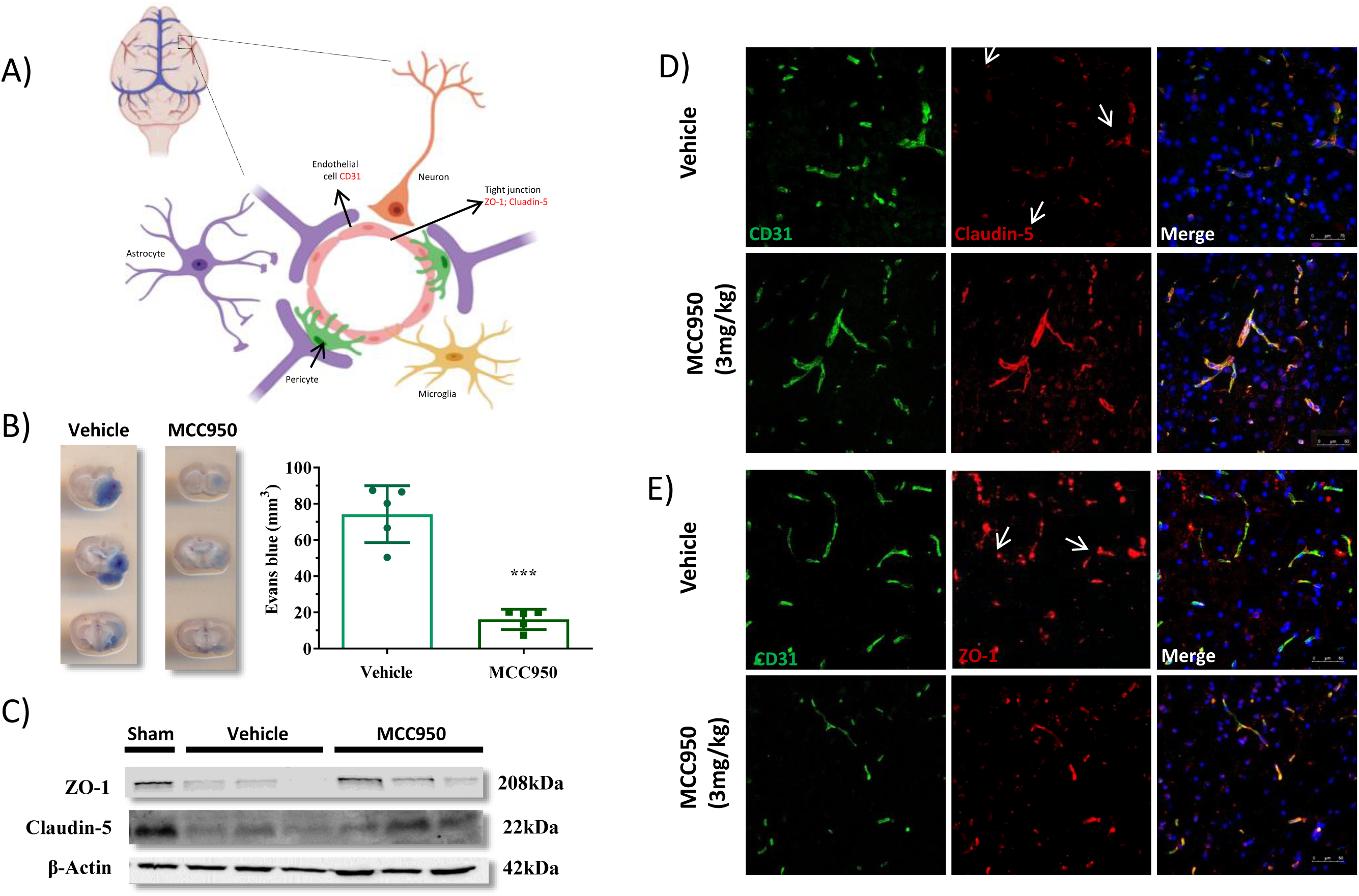
MCC950 (3mg/kg) treatment preserves blood-brain barrier (BBB) integrity after brain ischemia. (A) Principal cellular components of the neurovascular unit are represented in the scheme. BBB is a specialized boundary that separates the blood from the central nervous system containing TJ proteins such as Claudin-5 and ZO-1. (B) BBB permeability as assessed by Evans blue extravasation was preserved in treated animal with MCC950 3mg/kg but not in the vehicle-treated mice 24 hours after 1 hour of tMCAO. Complete set of brain slices from a representative animal are shown above the graph (Vehicle, n= 5; MCC950; n=5). (C) Representative western blot images of ZO-1 and claudin-5 of vessel enrichment 24 hours after stroke in vehicle-treated and MCC950-treated mice. β-actin was used as the loading control. Quantification of western immunoblotting for ZO-1 and claudin-5 in each group (n = 3–4/group). D) Representative confocal images of Claudin-5 and CD31 double immunostaining in the brains obtained after tMCAO in vehicle-treated and MCC950-treated mice. White arrows point the gaps junctions between adjacent tight junction proteins. Scale bar = 50µm. Data are normalized to sham. Data are mean ± SD. ***p<0.001 compared to vehicle mice.

Tight junctions (TJ) proteins form the initial barrier at the endothelial cells between blood and brain cells ^24^. In fact, TJ disruption is a major cause underlying the increased paracellular permeability of the BBB after ischemic stroke ^25^. To identify the role of NLRP3 inflammasome inhibition in TJ proteins degradation, i.e. Claudin-5 and Zonula Occludens-1 (ZO-1), we measured these proteins in enriched microvessels isolated from sham, ischemic and MCC950-treated animals (Fig. 4C). To validate the correct microvessel enrichment, we examined the levels of an astrocyte specific marker, GFAP, resulting in less GFAP in the microvessel enrichment than in the total brain tissue, demonstrating a successful microvessel enrichment (Fig. S1). Second, we analyzed TJs disruption by measuring the protein levels in cerebral microvessels enriched isolated from ischemic hemisphere of the different experimental groups. The analysis of cerebral microvessel enrichment showed a significant decrease of ZO-1 and Claudin-5 protein expression in stroked mice, which correspond to a disruption of BBB. Interestingly, MCC950 (3 mg/kg) treatment partially restored TJs proteins in enriched cerebral blood microvessels (Fig 4C). Additionally, we studied Claudin-5 and ZO-1 expression counterstained with the endothelial marker CD31 by immunofluorescence. We identified a gap formation at the endothelial cell membrane of blood vessel in ischemic animals caused by the rearrangement of Claudin-5 and ZO-1. Surprisingly, the abundance of gap formation in MCC950-treated mice was significantly decreased (Fig.4D, E). Thus, NLRP3 inhibition remains as a crucial player in BBB stability after brain ischemia by reducing the loss of the TJ proteins Claudin-5 and ZO-1, gap formation and, ultimately BBB disruption.

### NLRP3 inflammasome inhibition reduces the expression of MMP9 and the different endothelial adhesion molecules (VCAM, ICAM-1) after brain ischemia

In the ischemic scenario, inflammatory cells release cytotoxic agents, i.e. matrix metalloproteinases (MMPs), which lead to reversible degradation of TJs during early stages after the onset of ischemia ^26^. MCC950 treatment reduced MMP9 expression in the brain of stroked mice 24h after MCAO, both at mRNA and total protein levels (Fig. 5A). These data correlate with the reduction of Evans blue extravasation and TJs loss observed after brain ischemia. Endothelial adhesion molecules (VCAM, ICAM-1) mediate interaction between leukocytes and endothelial cells during the inflammatory response ^27^. To investigate the role of NLRP3 inflammasome inhibition in endothelial cells adhesion molecules, we assessed ICAM-1 and VCAM gene expression in enriched microvessels. We observed an increase of ICAM and VCAM mRNA levels in stroked animals after 24 h, which was significantly reduced with MCC950 (3 mg/kg) (Fig. 5B). Altogether, our results suggest that MCC950 can promote the protection of the neurovascular unit, preventing BBB leakage by reducing MMP9 total levels, TJs loss and the production and release of CCL2 and endothelial adhesion molecules (ICAM-1 and VCAM) after brain ischemia.

**Figure 5.**
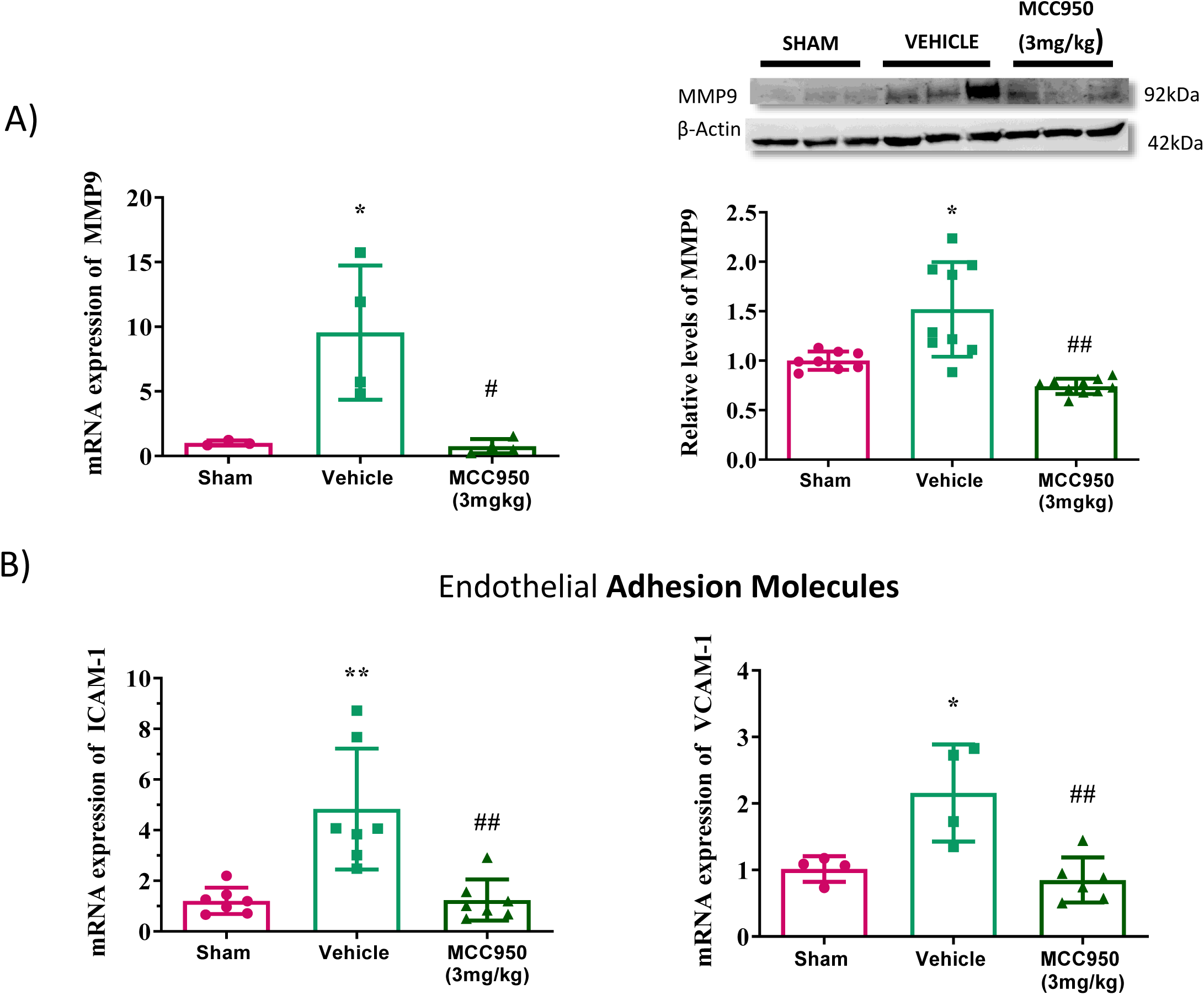
Inflammasome NLRP3 inhibition reduces the expression of different endothelial adhesion molecules (VCAM, ICAM), MMP9 and CCL2 in brain vessel enrichment after stroke. (A) Representative western blot images of MMP9 24 hours after 1 hour of stroke in mice treated with the vehicle and MCC950 (3mg/kg). β-actin was used as the loading control. Quantification of western immunoblotting for MMP9 in each group (n = 9–10/group). Data are normalized to sham. MMP9 mRNA levels of brain vessel enrichment after 24 hours of cerebral ischemia vehicle-treated and MCC950-treated mice. Data are normalized to sham. (B) Endothelial adhesion molecules (VCAM, ICAM) mRNA levels in brain vessel enrichment after 24 hours cerebral ischemia mice treated with MCC950 and vehicle. Data are mean ± SD. *p<0.05; **p<0.01compared to sham mice; #p<0.05 and ##p<0.01 compared to vehicle mice.

### Peripheral immune cells are increased in the ipsilateral hemisphere 24 hour post-stroke

Peripheral immune cells access to the brain is prevented by the BBB, but under pathological conditions, such as stroke, they can adhere to the activated post-capillary venules and infiltrate the brain parenchyma across the BBB ^28^. For the quantification of the peripheral immune cells infiltration into the cerebral parenchyma we performed flow cytometry studies using brain tissue samples of sham-operated mice, vehicle and MCC950 treated mice at 24 hours of reperfusion (Fig. 6A, B). FACS analysis of the number of microglial cells (CD45int CD11b+) in the ipsilateral hemisphere showed no significant difference across the groups (Fig. 6C). However, the number of total leukocytes (CD45HI) and myeloid cells (CD45HI CD11b+) were increased in the ipsilateral hemisphere of vehicle-treated mice compared to sham-operated mice 24 h after brain ischemia (Fig. 6D, E). Interestingly, treated mice showed a decrease in the number of leukocytes and myeloid cells at 24 hours of reperfusion (Fig. 6D, E). Recent experimental studies show that neutrophils migrate to cerebral ischemic regions during the first few hours after the onset of ischemia ^29^. Finally, we measured the number of polymorphonuclear cells (PMN) such as neutrophils at 24 h after brain ischemia. We observed an increase of PMN (CD11b+ Ly6G+) in the ipsilateral hemisphere at 24 h post-reperfusion in vehicle-treated mice compared to sham-operated mice. Similarly, MCC950-treated mice showed decreased numbers of PMN (Fig. 6F). To confirm these findings, we performed immunofluorescence labelling of the PMN marker Ly6G. Likewise, these results proved fewer Ly6G positive cells in the cerebral ischemic regions in MCC950-treated mice compared to vehicle-treated mice (Fig. 6G). Altogether, these results demonstrated that 1 h of MCAO followed by 24 h of reperfusion induced the recruitment of peripheral immune cells into cerebral ischemic regions and that MCC950 (3mg/kg) treatment 1 h after reperfusion had the potential to reduce recruitment.

**Figure 6.**
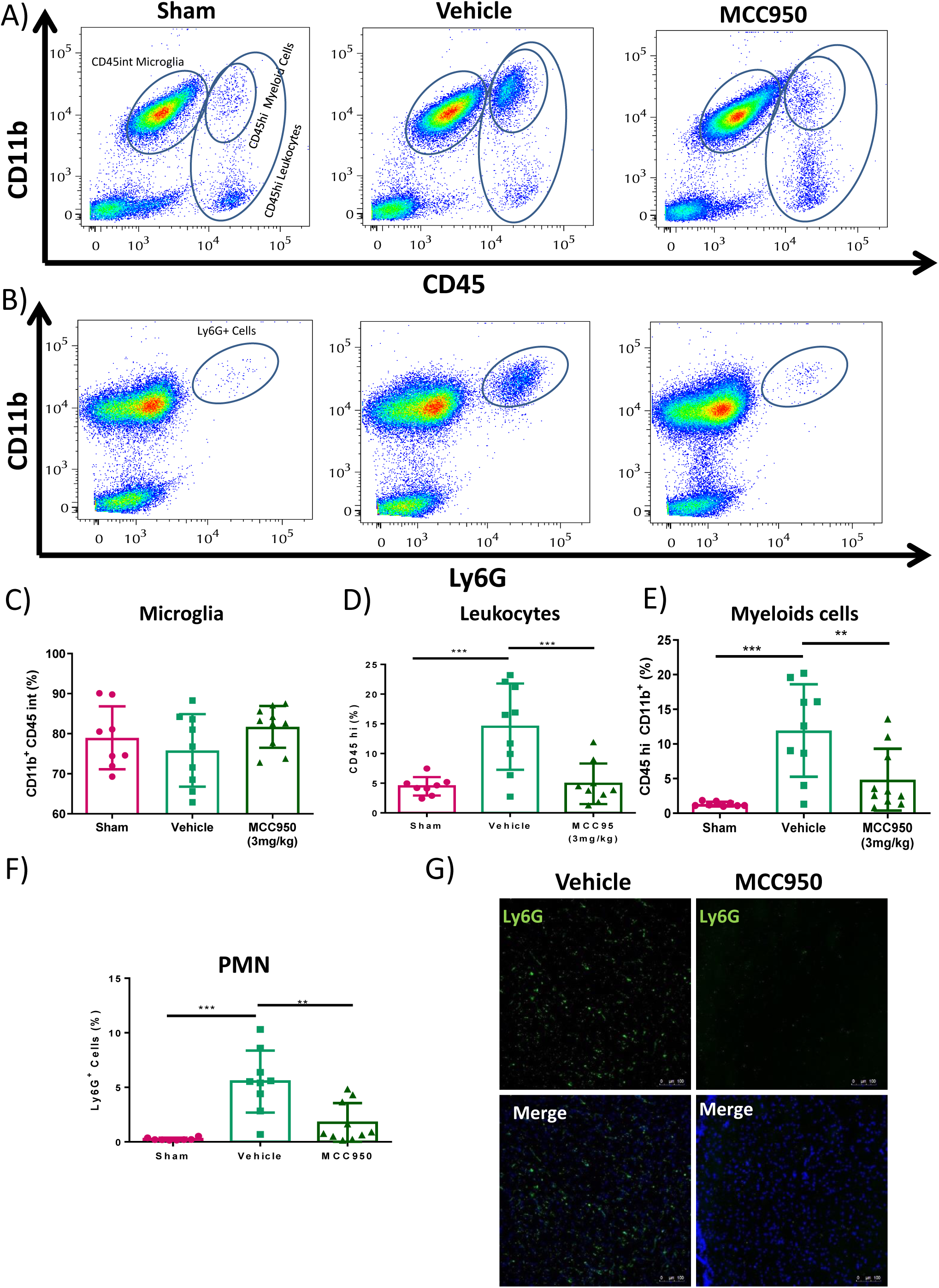
Peripheral immune cells infiltration into the brain 24 hours after brain ischemia. Immune cells were isolated from the ipsilateral hemisphere and assessed via flow cytometry 24 hours post-stroke. A) Representative flow data plots depicting the gating strategy used to identify brain-resident CD45INT microglia, CD45HI leukocytes, CD45HI CD11b+myeloid cells and B) CD11b+Ly6G+ polymorphonuclear cells at 24 hours after tMCAO in mice treated with vehicle and MCC950 (3mg/kg). (C) Quantification of brain microglia, leukocytes (D), myeloid cells (E) and polymorphonuclear cells (F) 24 hours post-stroke in vehicle-treated MCC950-treated mice, expressed as a percentage over total alive cells. (G) Representative confocal images of Ly6G in the brains obtained from tMCAO mice treated with vehicle and MCC950. Scale bar = 100µm. For all experiments, sham (n=8), vehicle (n=9) and MCC950 (n=10). Data are normalized to sham. Data are mean ± SD. **p<0.01; ***p<0.001 compared to vehicle mice.

## Discussion

Here, we report a time-dependent dual effect of NLRP3, the canonical sensor of sterile injury, in the acute phase of ischemic stroke. We have shown that ischemic brain injury was not reduced when NLRP3 was inhibited or absent before the ischemic onset, either by specific inhibition with MCC950 in pre-treated mice or in NLRP3 KO mice approach. However, post-reperfusion treatment with MCC950 (from 1 to 2 h after ischemic onset) had a strong protective effect of the neurovascular unit. Post-reperfusion inhibition of NLRP3 led to a reduction of “cytokine storm”, decreasing the levels of circulating CCL2 and MMP9, and hence protecting the BBB and TJs disruption triggered by cerebral ischemia and reduces the number of infiltrating neutrophils in brain parenchyma. These data provide evidence that NLRP3 has two roles in the acute phase of ischemic stroke. Hence, pharmacological inhibition of NLRP3 could break the inflammatory response produced after the ischemic onset, leading to a potent protective effect of the neurovascular unit.

In the last few years, numerous studies have been conducted to explore the participation of NLRP3 in sterile brain injury and diseases, such as cerebrovascular diseases (ischemic and hemorrhagic stroke), neurodegenerative diseases (Alzheimer’s, Huntington’s, and Parkinson’s diseases), multiple sclerosis, depression as well as other CNS disorders. Actually, there is some controversy over NLRP3 involvement in stroke. On the one hand, previous studies have reported that NLRP3 inflammasome inhibition or absence plays a vital role in the onset and progression of stroke, reducing the brain damage and protecting the brain ^12,13^, whereas on the other hand, it has been reported that mice deficient in NLRP3 had no reduction in brain injury ^10,11.^

In this study, we try to clarify the double-edge sword of NLRP3 inflammasome activation. First, we corroborate the data obtained with NLRP3 KO mice, where we show no reduction in ischemic damage; however, there is an increase in the mortality rate of this group, which indicates the importance of NLRP3 activation in the ischemia onset. These data lead us to distinguish between NLRP3 activation prior (NLRP3 KO and pre-treatment with MCC950) and after the ischemia onset (post-reperfusion treatment, 1 and 2 h). Taking this distinction into account, we obtained the same results; in the first group (prior to ischemia onset), we showed a lack of reduction in the ischemic damage and an increase in the mortality rate; in the second group (after the ischemia onset), we obtained a significant protection against brain ischemia with no change in the mortality rate compared to ischemic animals. These data lead us to rise the hypothesis that NLRP3 is important following brain ischemia and needs to be activated right after the ischemia onset to control the damage, but this activation needs to be controlled in the time-window of 1 h after the ischemic onset to prevent hyperinflammation due to BBB disruption.

IL-1β is one of the most extensively studied cytokines in the context of neuroinflammation. In the context of stroke, it has been demonstrated that IL-1β KO mice show smaller infarcts than WT and IL-1β administration worsens the outcome of ischemic rats ^30,31^. Therefore, the participation of IL-1β in stroke is clear. However, which inflammasome is involved and what is the participation of central and/or peripheral IL-1β is still under debate. In this work, we elucidated the participation of NLRP3 in acute brain injury, clearly differentiating its participation before and after the ischemic onset. We therefore added new evidence of NLRP3 taking part in brain ischemia, so it can be included to the list of inflammasomes that have been demonstrated to participate in brain ischemia, such as NLRC4 and AIM2 ^10^. However, more information is needed to discriminate between the effect of central and peripheral IL-1β. It has been recently shown that systemic inflammation, either produced by direct IL-1β injection or LPS-induced IL-1β release, drives to BBB disruption and brain inflammation ^32–34^. Unilateral overexpression of IL-1β in mouse brain leads to infiltration of immune cells into the hippocampus and BBB damage ^35^. Thus, further studies are required to figure out this unsolved question.

We have previously shown that toll-like receptor 4 (TLR4) blockage has a strong protective effect in different models of brain ischemia, including in humans ^14^, and this protective effect is mainly based on the control of the microglial inflammatory state. In this study, we sought to deepen into the comprehensive study of the fine-tune control of inflammation, by inhibiting NLRP3 activation to control innate immunity over-activation, and what we observed is that the specific inhibition of NLRP3 had a potent protective effect on the BBB in brain ischemia/reperfusion. The opening or rupture of BBB has been shown to be biphasic ^36^. In the first phase, ischemia-reperfusion induces subtle cytoskeletal changes, essentially junctional proteins redistribution in the microvascular endothelial cells, followed by MMP9 increases, and enzymatic cleavage of TJ proteins after inflammatory cell recruitment mediated by chemokine liberation (CCL2/MCP-1, CCL3, CXCL12/SDF-1), leading to the irreversible BBB breakdown and to secondary expansion of tissue injury ^37^. Here we show that the potent protective effect of MCC950 relays on the stabilization of TJs at endothelial level in this secondary phase. Hence, MCC950 reduces: i) MMP9 expression (Fig 5), ii) CCL2 levels in brain and in serum (Fig 3), iii) expression of the endothelial adhesion molecules ICAM-1 and VCAM-1 (Fig 5), iv) inflammatory cells recruitment, mainly polymorphonuclear cells (Fig 6) and, finally, v) the enzymatic cleavage of TJ proteins, leading to strong BBB protection (Fig 4), reduction of infarct volume and neurological outcome (Fig 1). All the effects were mediated by reduction of NLRP3-induced IL-1β liberation.

## Summary

In summary, our data provide new pieces of evidence showing the crucial role of NLRP3 activation in the pathophysiology of brain injuries. We have shown that NLRP3 activation is necessary for animal survival but there is a therapeutic window after the ischemic onset in which NLRP3 can be blocked to obtain a strong protective effect. Overall, our data show that improvements on BBB integrity by NLRP3 inhibition during the first few hours following an ischemic insult could be used as a target for therapeutical treatment that may limit neurological damage induced by pro-inflammatory cytokine/chemokine release and inflammatory cell infiltration.

## Non-standard Abbreviations and Acronyms

DAMPs: damage-associated molecular patterns
PAMPs: pathogen-associated molecular patterns
NLRC4: NACHT, LRR, CARD domain-containing protein 4
NLRP3: NACHT, LRR and PYD domains-containing protein 3
AIM2: Absent In Melanoma 2

## Acknowledgments

We are grateful with P. Pelegrín (IMIB-Arrixaca, Spain) for the *Nlrp3*–*/*– mice. We also thank Instituto/Fundación Teófilo Hernando for its continued support.

## Sources of Funding

This work was supported by grants from Fondo de Investigaciones Sanitarias (FIS) (ISCIII/FEDER) (Programa Miguel Servet CP14/00008; CPII19/00005; PI16/00735; PI19/00082) and Fundación Mutua Madrileña to JE. Kootstra Talented Fellowship (UM, The Netherlands) to AC. Grants from the Spanish Government (co-funded by European Regional Development Fund, ERDF/FEDER); PI16/02166 and Red Temática de Excelencia en Investigación en Hipoxia (SAF 2017-90794-REDT) to MJC.

## Disclosures

The authors have declared that no conflict of interest exists.

## Notes

### Competing Interest Statement

The authors have declared no competing interest.

