## Supplemental files for "Time-dependent dual effect of NLRP3 inflammasome in brain ischemia"

Short title: Key role of NLRP3 inflammasome in brain ischemia.

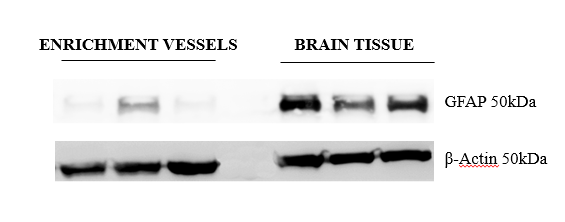

**Fig. S1. Brain vessel enrichment**. A) GFAP was used as a control protein to vessels enrichment from brain total tissue.

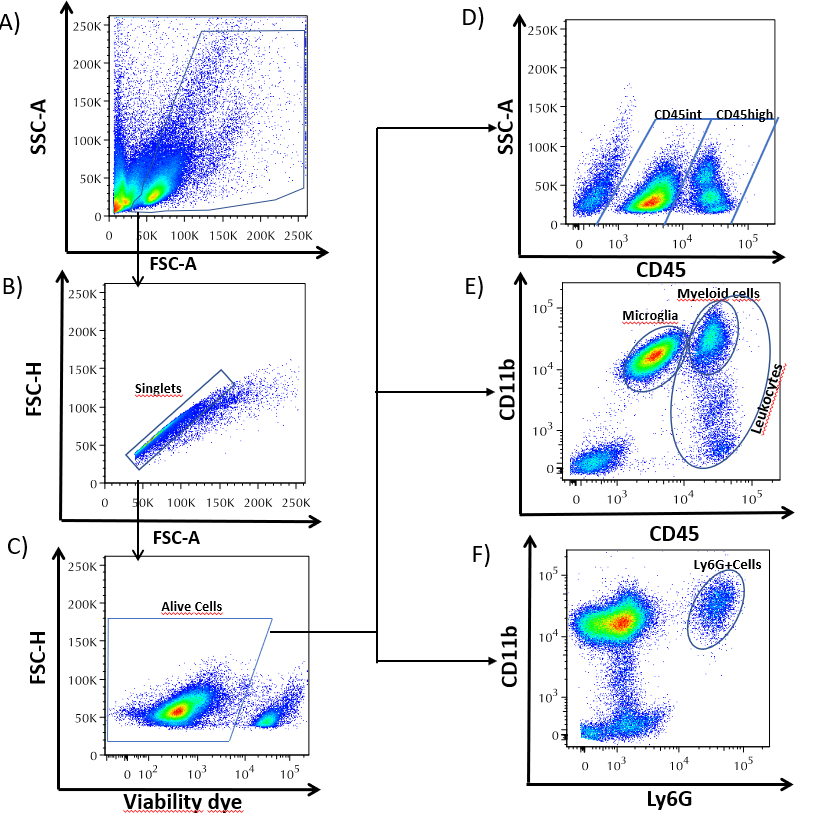
Fig. S2. Representative gating strategy for flow cytometric analysis of immune cells in the brain, at 24 hours post-stroke. (A) Cell populations are first identified by side and forward scatter (SSC and FSC). (b) Singlets were gated on FSC-A versus FSC-H (C) and live cells were gated based on Ghost viability dye staining. (D) CD45+ immune cells were gated based on CD45 staining versus SSC-A. (E) Cell surface antigens CD11b and CD45 were used to identify myeloid cells and microglia. (F) CD11b versus Ly6G were used to identify PMN cells.

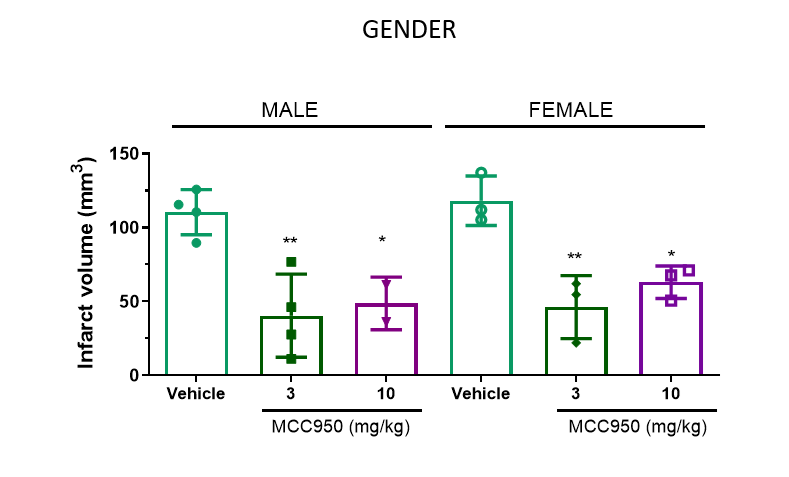
**Fig. S3. Effect of NLPR3 Inflammasome in cerebral ischemia 24 hours after 1 hour of stroke in male and female mice.** (A) Representative images and quantification of infarct volume by TTC staining at 24hours after 1 hour of ischemia in male and female mice treated with vehicle (male n=4; female n= 3) and MCC950 at 3mg/kg (male n=4; female N=3) and 10 mg/kg (male n= 2; female n=3). Data are mean ± SD. *p < 0.05; ** p < 0.001.

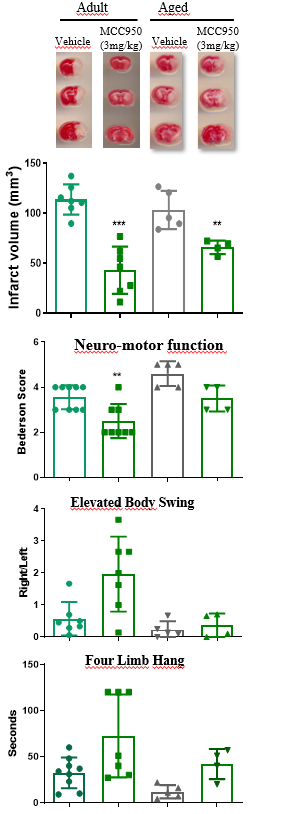
**Fig. S4. Effect of NLPR3 Inflammasome in cerebral ischemia 24 hours after 1 hour of stroke in aged mice.** (A) Representative images and quantification of infarct volume by TTC staining at 24hours after 1 hour of ischemia in aged mice treated with vehicle (n=5) and MCC950 3mg/kg (n=4). (B) Quantification of Bederson score 24 hours after 1 hour of ischemia in adult mice versus aged mice treated with Vehicle (n=9; n=5) and MCC950 3mg/kg (n=8; n=4), respectively. (C) Quantification of elevated body swing test 24 hours after 1 hour of ischemia in adult mice versus aged mice treated with Vehicle (n=7; n=5) and MCC950 3mg/kg (n=7; n=4), respectively. (D) Quantification of four limb hanging test 24 hours after 1 hour of ischemia in adult mice versus aged mice treated with Vehicle (n=7; n=5) and MCC950 3mg/kg (n=7; n=4), respectively. Data are mean ± SD. *p < 0.05; ** p < 0.001.

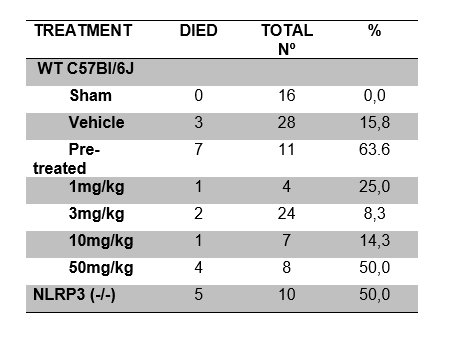
Table S1. Mortality rate. Percentage of deaths in WT C57Bl/6J treated with MCC950 at different doses and NLRP3(-/-) mice after 1 hour of tMCAO.
